# A Purpose-Built Open Source Liquid Handler for Industry-Class Automated Experiments

**DOI:** 10.64898/2026.03.02.709168

**Authors:** Stefan Golas, Bryant Gill, Kyle Wardlow, Aidan Baydush, Johannes Linzbach, Emma Chory

## Abstract

The expanding scope of laboratory automation increasingly demands systems that can be tailored to specific experimental constraints, including footprint, timing, cost, and control. While open-source software has improved protocol flexibility, liquid-handling hardware itself remains largely closed, limiting the ability of academic and startup laboratories to build instruments around biological requirements rather than vendor defaults. Here, we present a fully open-source, purpose-built liquid-handling robot assembled from commercially available components and developed entirely in a research setting. The platform integrates open hardware, electronics, and a Python-based control stack compatible with PyLabRobot, exposing low-level motion dynamics and liquid-handling behaviors directly to experiment code. We validate the system using a high-throughput turbidostat workflow that requires rapid, closed-loop measurement and actuation to maintain microbial cultures at defined density setpoints. The robot sustains stable steady-state growth across approximately 200 cultures with heterogeneous growth dynamics. A replica build completed by two lab members in approximately one week confirms that the platform can be reproduced from its bill of materials and assembly guide. Its compact footprint and use of off-the-shelf components make it suitable for rapid, parallel deployment in settings such as public health emergencies or by distributed laboratories. Together, these results demonstrate that industry-class liquid handlers can be custom-built for specific experimental goals, establishing a blueprint for open, purpose-driven hardware development across research and industrial automation contexts.

Open Liquid Handler (OLH) Design Goals.
**Left**: Design goals for a purpose-built platform for time-sensitive, closed-loop biological workflows, emphasizing high-accuracy dosing (low variability liquid handling), rapid integrated measurement (plate deck and isolated workspace), customizable deck and peripheral options, compact footprint with high throughput, containment via an enclosed wet workspace for biosafety and sterility, and a replicable build using off-the-shelf OEM components with open design files. **Right:** Open Liquid Handler design and physical implementation, with aerial and front views highlighting the enclosed cabinet and the working envelope over a compact deck.

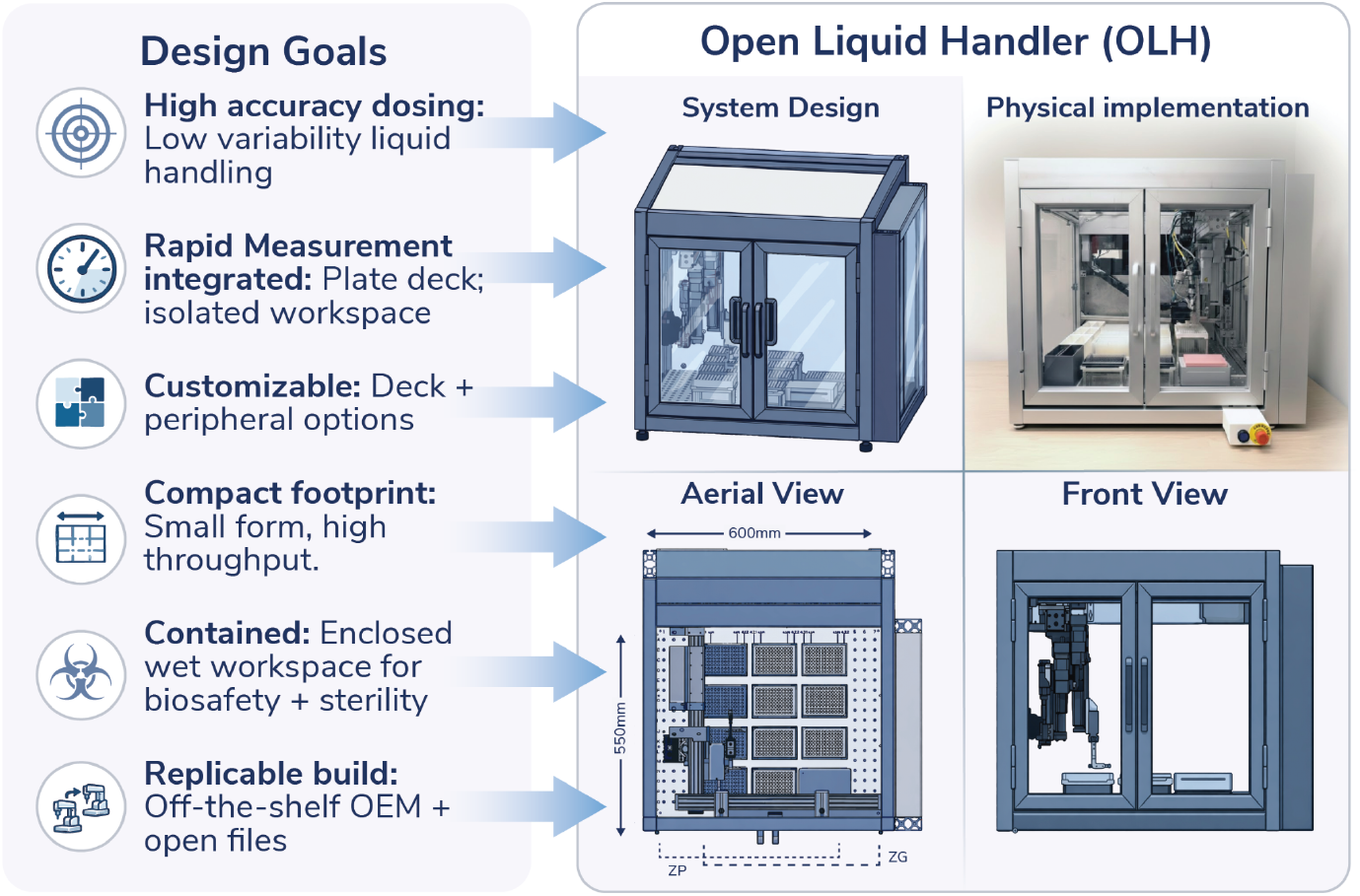

## Introduction

Automation has become central to improving reproducibility, experimental scale, and the collection of rich, time-resolved datasets. However, the intrinsic pace of many biological processes has shaped how automation is typically applied. As a result, most automated workflows are organized around static, batch-style operations such as sample preparation (PCR, diagnostic workflows^1^), *in vitro* end-point assays^2^, or media exchange at fixed intervals^3^, where timing flexibility is less critical. These processes assume relatively low time sensitivity, infrequent intervention, or can operate on predetermined schedules rather than requiring continuous responsiveness. As experimental designs evolve toward closed-loop operations, real-time sensing with decision-making, and adaptive or AI-guided optimization across many parallel conditions, the limits of batch assumptions begin to be stressed.

Traditional liquid-handling robots are powerful, reliable, and supported by mature hardware and software ecosystems, and have enabled substantial gains in throughput for standardized assays. These platforms are exceptionally suited for executing predefined protocols reproducibly at scale, for integrating common laboratory consumables and peripheral devices, and for deployment in regulated environments that require validated operation and traceable documentation (for example, GMP- or GLP-aligned workflows). However, time-critical workflows expose gaps in conventional liquid-handling systems, as most platforms are optimized for broad use, and are not readily reconfigurable for custom, needs-based, biology-timed experiments. Low-level behaviors such as gantry motion profiles, acceleration limits, pipetting speeds, dwell times, and task scheduling are typically fixed or remain indirectly accessible through proprietary interfaces. This limits the achievable control bandwidth and the range of closed-loop experiments that can be implemented with industrial-grade systems.

To meet this need, DIY automation efforts have lowered the barrier to building, modifying, and sharing laboratory instruments, motivated both by frugal-science goals^4–6^ and by the need to prototype capabilities outside vendor ecosystems. On the software side, open control layers such as PyHamilton^7^ and PyLabRobot^8^ have shown that programmatic interfaces can unlock complex behaviors on existing liquid handlers and make protocols more portable by decoupling experimental logic from proprietary environments. In parallel, open hardware platforms have demonstrated that functional laboratory instruments can be assembled from accessible components with shared designs, including low-cost DIY liquid-handling robots for education and early-stage research^9^, programmable chemical synthesis platforms^10^, as well as commercially available open platforms^11^. Related “frugal science” efforts have similarly shown that rigorous scientific tools can be designed for accessibility and broad dissemination^12^, and several of these projects have been translated into companies or community-deployed platforms, illustrating how academic prototypes can evolve into standardized, transferable workflows. However, most open systems intentionally prioritize education or a narrow experimental scope, leaving open questions about how to achieve the mechanical capability, reliability engineering, and system integration expected of industry-grade instruments used in regulated, manufacturing-adjacent, or even patient-facing settings.

Biological reactors are a useful stress test for data-responsive automation because they require sustained operation under application-specific constraints. For nearly a century, process control engineering has enabled continuous sensing and feedback control in manufacturing-scale bioreactors by regulating variables such as pH, dissolved oxygen, temperature, agitation, and feed profiles^13^. At the research scale, bioreactor platforms such as eVOLVER^14^ and Chi.Bio^15,16^ have expanded adoption of feedback-controlled continuous culture systems. Additionally, we previously adapted a commercial liquid-handling platform for high-throughput, feedback-controlled microbial culture by integrating plate reader measurements with real-time, Python-based decision making^7^. However, as data-responsive workflows expand in scale and diversity, there is a continued need for industry-grade liquid handlers that can be purpose-built for time-critical workflows while remaining affordable and straightforward to replicate across laboratories and applications.

To our knowledge, no industry-class liquid handler has been made available as a fully replicable, open build. As a result, groups have limited ability to make workflow-specific mechanical tradeoffs when writing automation protocols. As an example, experiments that maintain cell populations at a defined density setpoint over time would be widely useful in research and biomanufacturing because they decouple cellular physiology from batch growth effects. In academic settings, continuous-culture approaches support studies of gene regulation, metabolism, stress responses, and evolutionary dynamics across diverse microbes; in applied settings, they underpin strain and process development for fermentation and cell-based production. When run in parallel, these methods enable systematic studies across strains, environments, and perturbations, from screening and characterization to long-term evolution experiments. However, simultaneously maintaining many cultures at a density setpoint creates practical engineering constraints. Traditional turbidostat bioreactors typically require dedicated pumps and sensors for each individual reactor, limiting scalability. While our existing liquid-handling robot approaches can multiplex measurement and dilutions across many cultures^7^, our current implementations require a large footprint and, in time-critical regimes, can be limited by dilution cycle time, making it difficult to maintain setpoints for rapid-growing strains or abrupt condition shifts. We therefore selected turbidostat-style control as a stringent demonstration case for a purpose-built, open liquid-handling (OLH) platform.

In this work, we provide a fully buildable, open-source liquid-handling robot and validate it on closed-loop, turbidostat-style culture maintenance at scale. The platform is assembled from commercially available motion and fluidic components, integrates eight independent pipetting channels with an on-board gripper in a compact ∼600 mm x 600 mm footprint, and achieves functionality comparable to industry-grade systems using parts that can be purchased for under $40,000. At a high level, the robot is controlled through a Python stack, exposing kinematic and liquid-handling parameters (including motion speeds, accelerations, and pipetting behaviors) directly to experiment code and enabling integration with on-deck peripherals such as plate readers, shakers, peristaltic pumps, and a custom 3D-printed tip-washing module. We demonstrate the system using a demanding turbidostat workflow that couples frequent density measurements to automated dilution, tip sterilization, growth-rate estimation across diverse conditions, and maintenance of user-defined setpoints across ∼200 independent cultures. Beyond this demonstration, the same implementation supports applications ranging from culture condition optimization for biomanufacturing, to growth phase-specific antibiotic response profiling by perturbing cultures at defined densities, to strain phenotyping and fingerprinting in constrained clinical lab spaces, and workflows requiring compact integration with anaerobic enclosures. More broadly, by making both the instrument, its assembly, and its control stack fully open, this platform will allow automation itself to be treated as a modifiable research substrate, supporting extension to new end effectors, peripherals, and closed-loop, data-driven experimentation, including self-driving labs and AI-guided experimental design.

## Results

### Design Goals and System Architecture

Many biological processes that could benefit most from automation are not batch assays, rather they are time-sensitive, living processes that require continuous sensing and intervention. In turbidostat-style continuous culture, each population must be measured and diluted on a cadence set by its growth rate in order to maintain a defined physiological state. When growth is fast, latency in the liquid-handling loop degrades density control of bacterial cultures, increases variability across replicates, introduces biological artifacts such as biofilming, and can destabilize long-term experiments, especially those that shift their metabolic states between exponential or stationary phase growth. These constraints become more severe when scaling to a hundred or more parallel cultures, where per-culture service time sets an upper bound on throughput and on the fastest growth dynamics that can be stabilized. These same constraints appear in bioprocess development workflows that require steady-state growth to compare media, induction timing, or strain variants under controlled conditions. Thus, the overall objective was to develop a purpose-built automation platform that can execute time-sensitive, closed-loop biological workflows, sustaining tight control of culture state across many parallel populations with low-latency measurement, decision-making, intervention, logging, and cleaning, with both density and reporter-based readouts.

This objective motivates a single, primary performance goal: a short, repeatable control loop whose end-to-end latency is low enough to stabilize fast growth dynamics even when multiplexed across many cultures. In practice, this requires high motion throughput and minimal travel overhead so per-culture service time stays low as the experiment scales. It also requires stable, low-variability liquid handling across thousands of repeated interventions, including multi-channel transfers that can deliver different volumes across wells within a single operation, industry-grade quality controls, and reliable, automation-compatible measurements that do not become the bottleneck (e.g. frequent plate reader transfers). A second goal was containment, both for sterility and for environmental stability during long runs. Time-sensitive workflows often operate for days to weeks and can be sensitive to temperature, evaporation, and contamination. The system therefore requires a self-contained wet workspace that supports sustained operation and isolates cultures from the external lab environment while keeping high-voltage and motion-control components safely segregated. A third goal was open-source software control. Closed-loop workflows depend on real-time data acquisition, growth-rate estimation, and control decisions, and the implementation must be easy to modify as the biology and peripherals evolve. By implementing as much of the control architecture as possible through PyLabRobot, future changes are software-first: new devices, deck layouts, control logic, and workflows can be added within a consistent Python framework, while preserving protocol portability and maintaining compatibility with Python-driven ecosystems. Additionally, practical deployment constraints shaped the system-level design, including reproducibility across sites. As many microbial culture applications are inherently low marginal cost per condition (using mainly sugar, yeast extract, water, as primary consumables), automation hardware must not impose prohibitive capital or consumable costs. Similarly, space is often the limiting resource in academic, start-up and clinical laboratories, so the platform must perform within a compact footprint, rather than requiring a dedicated room-scale automation cell. Finally, the system must be adaptable and reproducible in other labs, which motivates a build strategy centered on commercially available components. This minimizes reliance on proprietary, single-source parts and lowers the barrier for groups that want to replicate the platform and then tailor it to their own workflows. To validate the reproducibility of the platform, a twin model was assembled to cross-validate protocols and assembly documentation **(Supp Fig. 1)**.

As such, the Open Liquid Handler (OLH) presented herein provides all of the above functional criteria using off-the-shelf Original Equipment Manufacturer (OEM) parts, with only a small number of components requiring custom machining or fabrication all of which are defined by public specifications and open-source design files so they can be readily fabricated, 3D printed, or produced in-house.

### Enclosure, cabinet, and deckware as a modular workspace

A fully enclosed layout was implemented to support containment and to ensure the motion system operates with consistent access constraints. The enclosure defines a repeatable coordinate frame for gantry alignment, provides fixed mounting points for deckware, and constrains splashes and aerosols to support safe unattended operation in a bio-processing environment that routinely encounters liquids and disinfectants **(Fig. 1A)**. The cabinet architecture physically separates the wet deck from the electronics cabinet, simplifying maintenance and reducing risk during iterative prototyping while preserving a stable mechanical reference between the motion stack and the working area. To make the workspace reconfigurable without redesigning the instrument, deckware was treated as a modular interface layer rather than a monolithic surface. Sheet-metal deck plates define the base coordinate reference and the global hole pattern used to locate access regions for Z end-of-arm tools and to register swappable fixtures across the deck **(Fig. 1B)**. At the fixture level, a standardized interface plate with defined slots and hole features (for example, M3 and M6) and key dimensions enables repeatable placement of labware nests, tip racks, reservoirs, and plumbing interfaces, supporting both standardized labware (for example, 96-well plates) and custom mounts fabricated by machining or 3D printing **(Fig. 1C)**.

**Figure 1).**
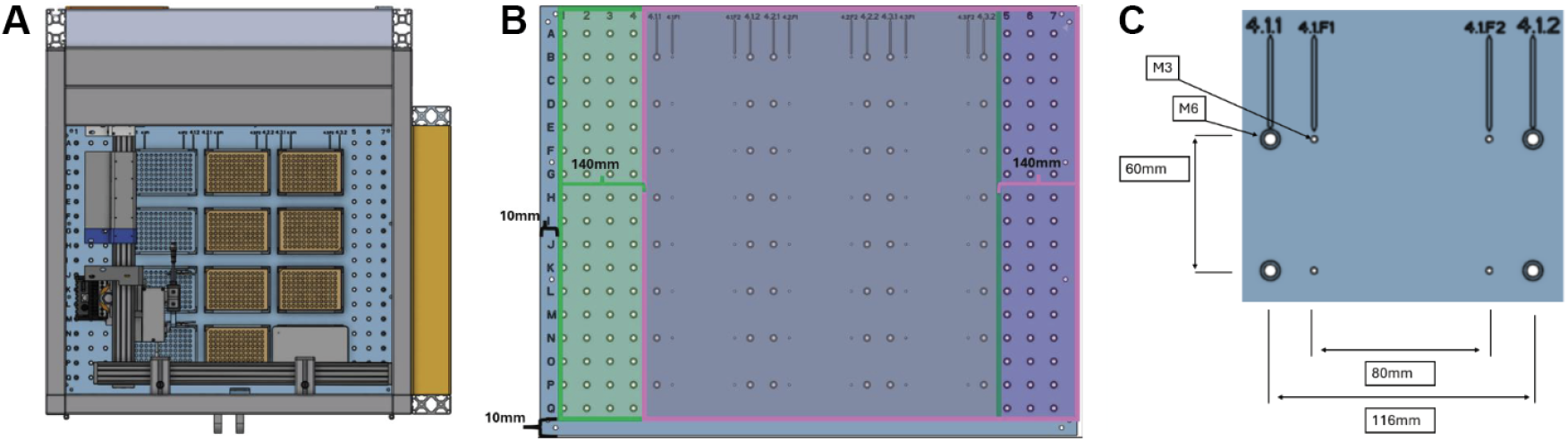
Enclosure, cabinet separation, and modular deckware for the Open Liquid Handler (OLH). **(A)** Top-down view of the enclosed instrument showing the wet-deck workspace and a physically separated side cabinet for electronics, providing a fixed coordinate frame and reducing exposure of electronics to liquids during iterative prototyping. **(B)** Sheet-metal deck plate layout and hole pattern that define the base coordinate reference and mounting regions for deckware, with the accessible regions for Z end-of-arm tools indicated: pipetting access 550 mm x 520 mm and gripper access 550 mm x 550 mm. **(C)** Example deckware interface plate detail with standardized slot and hole features (M3 and M6) and key dimensions, supporting swappable labware nests and fixtures with repeatable placement.

### Gantry motion system

The core motion system is a 3-DOF gantry that provides XY translation over the deck and Z translation for two independent end effectors. Along the X axis, a stiff guide rail establishes the reference geometry, while a linear actuator provides propulsion. Mechanical alignment at installation is critical: the guide rail is positioned relative to the cabinet frame and door-facing reference features, and the actuator is mounted so its motor and movement path remain co-linear with the guide rail across the full travel range **(Supp Fig. 2)**. Speed and repeatability of XY motion are central to the objective of this work. In turbidostat control, the scheduler revisits the same wells repeatedly with limited slack, and time spent moving between deck locations competes directly with time available for measurement, decision-making, and washing. Motion performance is therefore treated as a design constraint, and alignment checks are emphasized to preserve smooth high-speed motion across the full travel range. The X guide axis is placed near the door-side of the enclosure so that the gantry travel envelope aligns with the accessible deck region **(Fig. 2)**. Two placement checks reduce yaw and binding when coupling the Y axis to the X-axis slide plates: (i) lateral alignment of the guide axis relative to the cabinet corner caps and (ii) coplanarity of the guide rail end relative to the cabinet frame edge **(Fig. 2)**. The Y axis is assembled as a second linear rail that is orthogonally mounted to the X-axis slide plates via adapter kits **(Fig. 2)**. The Y-axis motor end is positioned toward the electrical cabinet region to shorten cable runs and reduce cable flexing within the wet workspace. The complete Y-to-X coupling geometry is summarized in **(Supp Fig. 2)**. Motor-to-actuator coupling is implemented using a stepper motor, a dedicated axial kit, and a centering procedure that aligns the coupling between the motor rotor and actuator interface before the motor is nested into the axial kit and the kit is locked to the actuator housing. Centering reduces eccentric loads on the motor bearings.

**Figure 2).**
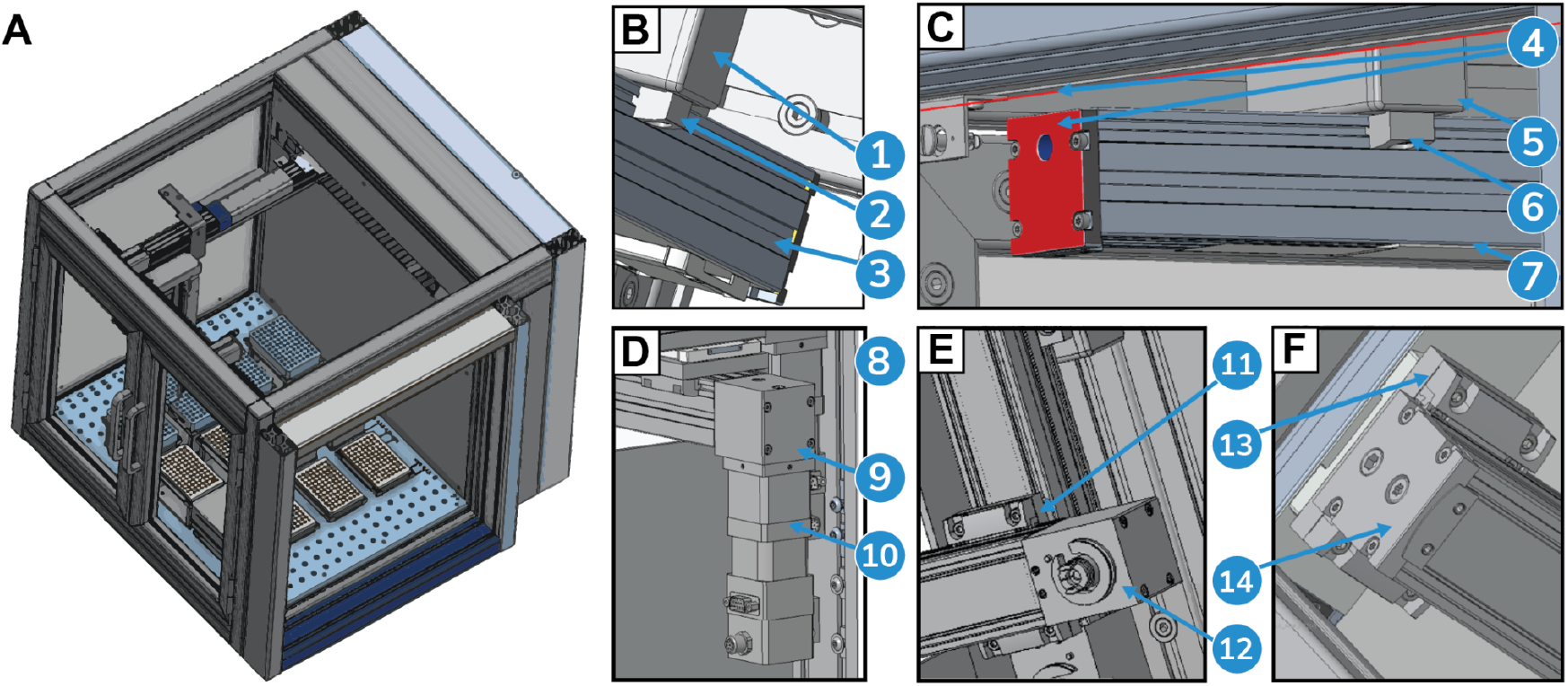
Gantry and motion stack of the liquid handling module. **(A)** Enclosed instrument showing the H-gantry spanning the deck and dual Z-axis toolheads (pipetting and gripper). **(B)** X-guide axis placement: X-axis guide rail [1] mounted to a Maytec aluminum profile via profile mounting hardware [2], integrated into the enclosure frame via a Maytec plate [3]. **(C)** X-guide rail alignment: guide rail face [4] aligned coplanar to the right-side cabinet frame edge (red reference line); rail mounts to a Maytec profile and plate [5] using profile mounting hardware [6], with the guide rail body shown in profile [7]. **(D)** Y-axis motor mounting: Y-axis ball-screw linear axis [8] mounted adjacent to the electrical cabinet; axial kit [9] couples the motor/drive train to the axis assembly [10]. **(E)** Y-to-X mounting (motor end): Y-axis motor end [11] interfaces to the X-axis drive via adapter kit mount [12]. **(F)** Y-to-X mounting (guide-rail side and non-motorized end): adapter kit [13] mounts to the X-axis guide rail, and the Y-axis non-motorized flat end [14] is captured on the carriage plate.

### Dual-Z plate handling and multi-channel pipetting

To support workflows that require both labware transport and liquid handling without tool changes, two independent Z slides are mounted to the Y-axis carriage. One Z axis is dedicated to plate handling via a parallel gripper, and the other Z axis is dedicated to liquid handling via a multi-channel pipetting head. This decoupling reduces scheduling constraints and allows a gripper to remain available for lid handling or plate movement while pipetting proceeds on a separate vertical stage. Plate handling and pipetting are separated into independent Z axes so the system transitions quickly between assay actions (lid handling, plate transfers) and control actions (dilution, sampling, reagent dosing). For closed-loop turbidostats, this reduces dead time between measurement and actuation and avoids forcing all operations through a single-end effector. Separation also improves fault isolation: a plate handling issue does not necessarily block pipetting, and vice versa. Each Z axis uses a ball screw-driven vertical slide architecture that supports compact integration on the Y-axis carriage while maintaining repeatable vertical motion **(Fig. 3A)**. The vertical stage provides a defined mechanical envelope for routing cables and, when needed, pneumatic lines to the end effectors while preserving a compact footprint. For plate handling, a parallel gripper is mounted to the Z slide through an adapter interface that sets the final tool geometry relative to the deck **(Supp Fig. 3)**. This modular interfacing approach supports reproducible instrumentation: commercially available parts implement the core kinematics and actuation, while thin adapter plates handle application-specific geometry and end-effector mounting. For liquid handling, a multi-channel pipetting head is mounted to the second Z axis through a corresponding adapter interface.

**Figure 3).**
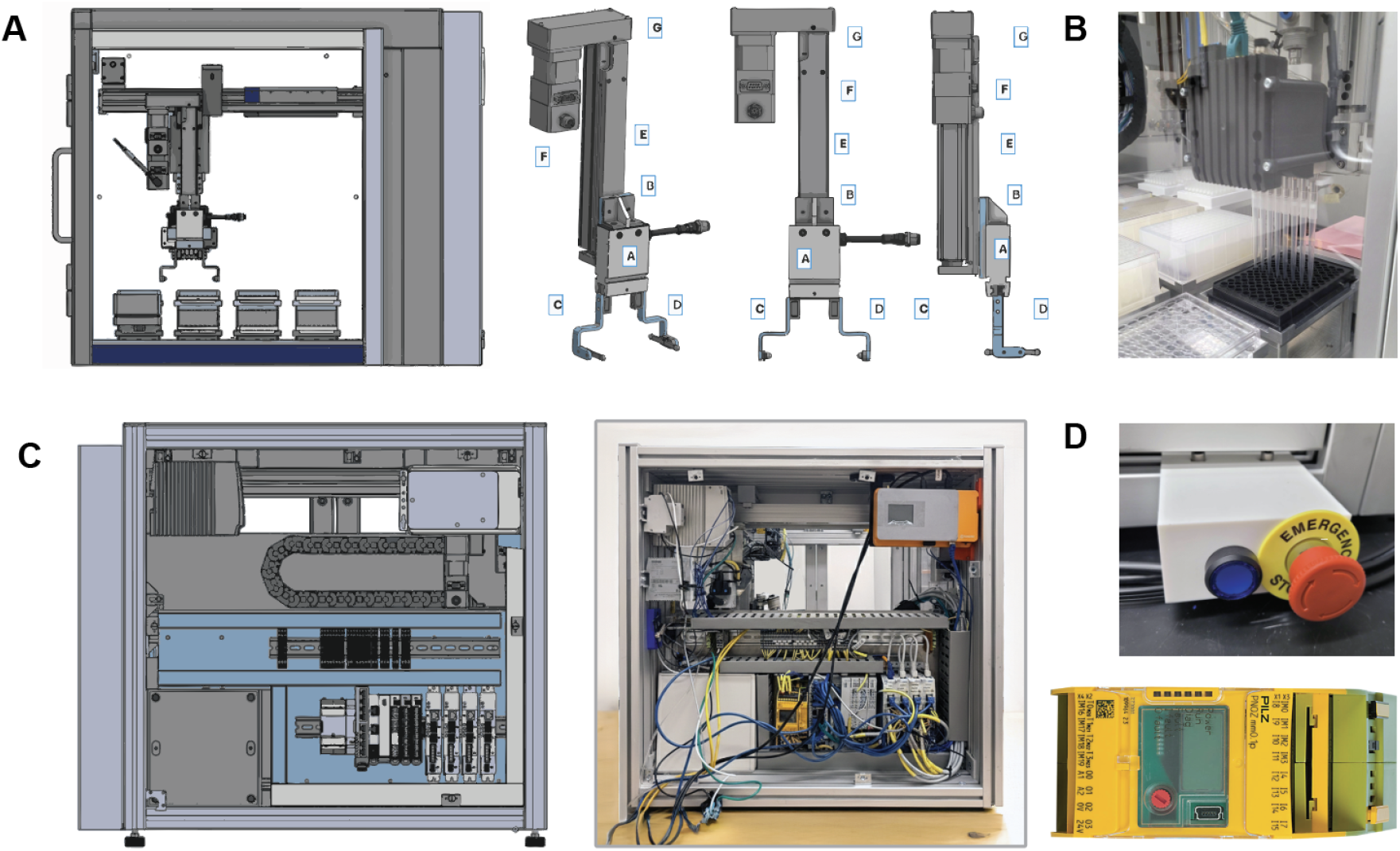
OLH Mechanical and Electrical Component overview. **(A)** Dual Z-axis end-effector stack for plate handling, showing the gripper-mounted Z slide and major mechanical subassemblies. **(B)** Liquid handling module in operation, showing the 8-channel pipetting head dispensing into a 96-well plate. **(C)** Electrical cabinet and controls, shown as a CAD layout and a representative build photo with DIN-rail mounted power distribution, remote I/O, and motion control components. **(D)** Safety interface hardware, including the enclosure emergency stop and reset controls used to immediately halt motion. *Image backgrounds were digitally removed using Adobe Photoshop Generative Fill (AI-assisted editing) to reduce visual clutter; no instrument features were added, removed, or altered*.

### Pneumatics and valve control

The pipetting stack relies on a combined pressure and vacuum supply with per-channel valve control to drive aspiration and dispense. The pneumatic architecture is organized as modular blocks for supply, regulation, distribution, and actuation, connected by standard tubing and push-in fittings **(Fig. 3B)**. This modularization supports maintainability by enabling rapid replacement of a leaking fitting, clogged valve, or failed regulator without disassembling the gantry. In turbidostat workflows, the same primitives repeat thousands of times: aspirate culture, dispense fresh media, mix, and periodically wash and sterilize. Pneumatic stability and valve timing therefore translate into biological stability because they influence delivered volumes, droplet formation, and the frequency of tip faults that can pause a run. Pneumatic and valve hardware is organized as explicit sub-assemblies so these parameters remain inspectable and serviceable during long experiments.

### Power distribution, remote I/O, and motion control electronics

The electrical cabinet is organized around DIN-rail mounting to support iterative revisions and serviceability **(Fig. 3C)**. Power distribution and wiring are structured to separate high-voltage mains handling from low-voltage logic and load wiring, with mains connection deferred until distribution blocks and downstream wiring are in place. The cabinet integrates an AC-to-DC supply for 24V logic power, terminal blocks for distribution and filtering, an Ethernet switch for network connectivity **(Supp Fig. 4)**, modular remote I/O for digital and IO-Link channels **(Supp Fig. 5)**, and stepper motor controllers for motion actuation **(Supp Fig. 6)**. The cabinet architecture emphasizes clean power distribution, clear service boundaries so modules can be replaced without rewiring the full system, and a wiring strategy that supports debugging under real operating conditions. Wiring is designed to provide distinct logic and load supplies where required, and to ensure correct supply pinning for motor controllers so undervoltage faults are avoided during commissioning **(Fig. 3C)**. The wiring architecture is captured in the schematic and cabinet layout documentation **(Supp Fig. 5), (Supp Fig. 6)**, which provides the authoritative connection map for assembly and service. Cable and line routing are treated as a core integration constraint in the enclosed, high-acceleration motion environment. Dedicated e-chain adapters and holders constrain bend radii and keep moving bundles within a repeatable envelope across the full travel range, reducing pinch points, connector torsion, and interference with deck hardware. Anchoring and bend orientation are defined relative to cabinet and frame references so routing remains consistent across builds. For long-term runs, cable routing must tolerate repeated high-acceleration motion without intermittent faults and must maintain separation between wet-deck hazards (splashes and disinfectants) and sensitive electronics. Cable management is therefore treated as part of the functional result: it enables stable long-duration operation while keeping troubleshooting straightforward when issues arise. Finally, an embedded compute module housed in the cabinet provides the instrument-resident server for gRPC communication with PyLabRobot, along with door-mounted service ports for AC mains entry and external data pass-through **(Supp Fig. 7A-B)**.

### Safety architecture

Safe operation is enforced through an enclosure interlock and emergency stop chain that removes motion power when the enclosure is opened and provides immediate shutdown via dedicated kill switches **(Fig. 3D)**. The safety chain uses standard components for enclosed motion systems, including a safety relay and magnetic door switches, and is integrated with the electrical cabinet power distribution so hazardous motion is inhibited during user access **(Fig. 3D)**. Safety validation checks are performed during commissioning to confirm that opening the enclosure reliably halts motion and that emergency stops produce an immediate system halt with a defined reset procedure. Because the OLH is intended to operate as an enclosed instrument during long, unattended runs, safety is implemented as part of the enclosure and cabinet architecture rather than as an external procedural control. Interlocks and emergency-stop circuitry establish a hard boundary between user access and motion while still supporting routine wet-lab operations such as loading plates, refilling media, and servicing waste without compromising containment.

### PyLabRobot implementation and software architecture

The Open Liquid Handler (OLH) is controlled in Python using PyLabRobot as the primary execution layer. The use of PyLabRobot as a programming interface is a decisively enabling factor in our robot’s advanced capabilities and low cost. Since PyLabRobot provides a generic and universally compatible interface to liquid-handling robots, we were not required to create an entire robot interface from scratch. Instead, we made use of PyLabRobot’s predefined generic hardware interface, deck layout management system, simulation module, and labware resource definitions to create an interface to our robot with minimal time investment or additional complexity. The use of PyLabRobot also facilitates accessibility due to its open-source license, cross-platform compatibility (e.g. to port scripts to other robots), and ease-of-use due to the use of the highly popular and readable Python language. Herein, PyLabRobot provides a consistent abstraction for deck resources, motion primitives, and device orchestration, which allows complex workflows to be expressed as composable, inspectable protocol code rather than as device-specific scripts **(Fig. 4)**. The software integrates four time-critical subsystems: (i) the liquid handling module for aspirate and dispense operations, (ii) a gripper device for plate transfer, (iii) an on-deck absorbance plate reader for automated measurements, and (iv) a washer and waste-handling module for cleaning and drain or bleach routines **(Supp Fig. 8)**. At the system level, the implementation emphasizes tight coordination between motion, peripheral I/O, and data capture. The codebase is organized as a small set of modules that separate core concerns: protocol orchestration, device interfaces, calibration utilities, and simulation support. This separation keeps hardware integration auditable and allows peripherals to be swapped, updated, or simplified without rewriting the core protocol logic. The repository also supports distinct execution modes that decouple experiment state from device access. Runs can be started fresh or resumed from disk, and protocols can be executed against real hardware or a simulation model for development and debugging. Persistent state and data outputs are treated as part of the method: each run writes a structured audit trail that records measurements, derived values, requested actions, executed commands, and error flags when present, enabling downstream analysis and reproducible re-execution.

**Figure 4).**
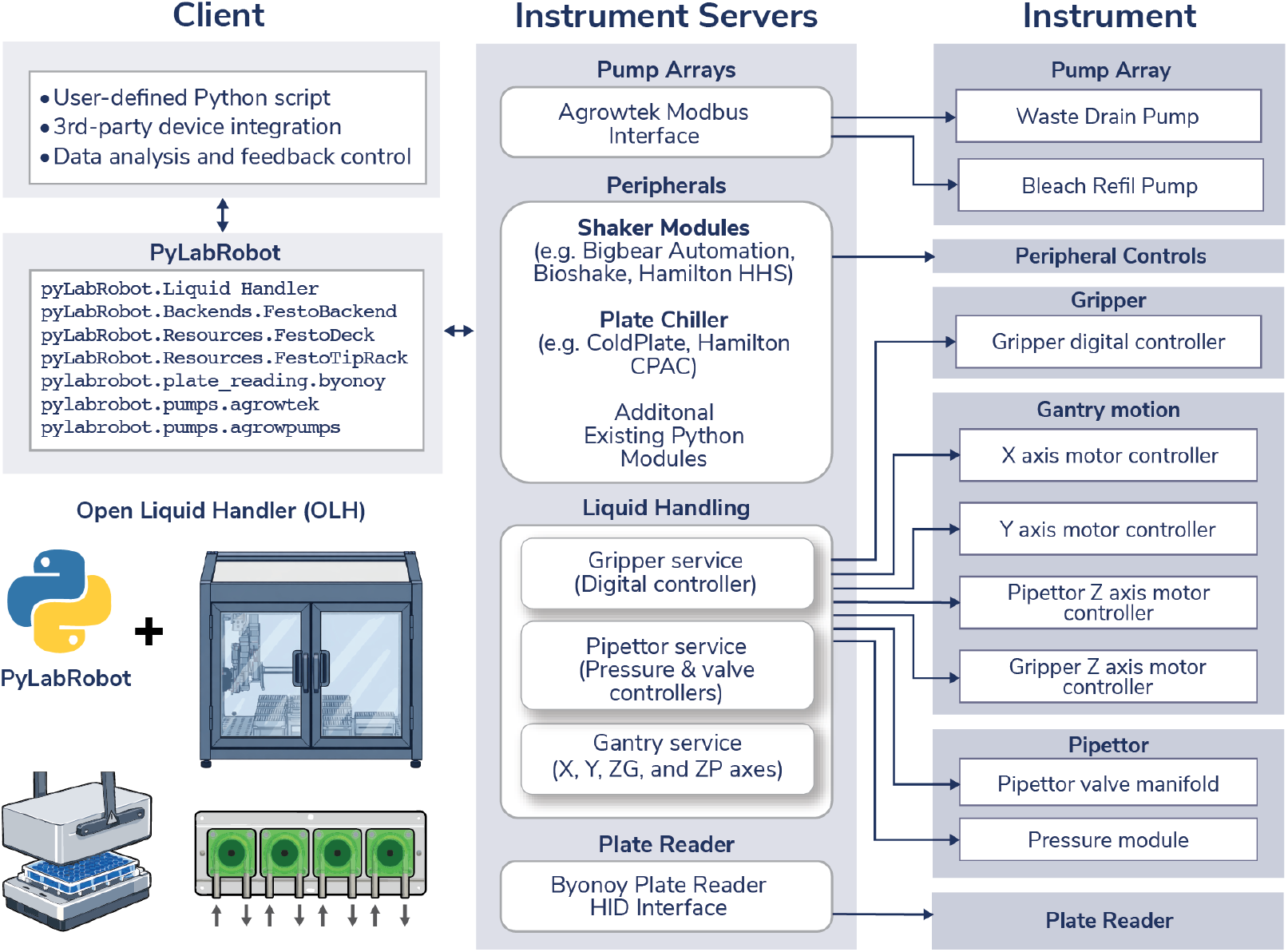
PyLabRobot software architecture for OLH control. A user-defined Python script interfaces with the OLH through PyLabRobot abstractions for the gantry, pipettor, gripper, plate reader, pumps, deck resources, and the OLH backend. PyLabRobot sends gRPC requests to an instrument-resident server. The instrument server processes incoming gRPC requests and implements the core control logic for coordinated movement (X, Y, and the gripper and pipettor Z axes), pipetting (pressure and valve control), and gripper actuation (digital control). The instrument server translates these requests into low-level commands for individual device controllers, including independent motor controllers for each axis and dedicated controllers for the pipettor pressure module and valve manifold and the gripper digital controller. Individual device controllers are interfaced through TCP Modbus interfaces, enabling modular peripheral integration while keeping protocol logic in a single Python layer.

### Pipetting Accuracy Calibration

Tip dosing accuracy is a critical requirement for closed-loop turbidostat control, where small, repeated volume allocations accumulate over hundreds of cycles. To quantify liquid-handling performance of the Open Liquid Handler (OLH), we generated an absorbance-based calibration curve using resazurin dilutions prepared manually with a calibrated P200 pipette. Serial dilutions were constructed across a defined dynamic range (20-200 uL) and measured on a BMG Vantastar F plate reader at 600 nm to establish a linear relationship between absorbance and volume. The calibration curve showed excellent linearity (R^2^ = 0.9984, n = 80) **(Fig. 5A)** and served as the ground-truth reference for subsequent accuracy benchmarking. We next used this calibration curve to validate robot-delivered volumes. The OLH executed identical dilution schemes, and measured absorbance values were converted back to effective volumes using the manually derived calibration function. Because the turbidostat workflow relies heavily on P1000-class transfers for media addition and dilution, we also directly compared OLH performance to manual multi-channel pipetting with a P1000 **(Fig. 5B)**. Manual dilutions were generated in parallel and evaluated using the same absorbance readout. Across the tested range, OLH-delivered volumes were significantly more accurate than manual multi-channel pipetting (mean absolute error 4.05 uL vs 7.17 uL; Welch’s t-test, t = -2.91, p < 0.005) **(Fig. 5C)**, with approximately half the error variance (ratio = 0.56), indicating superior precision under application-relevant conditions. Finally, we evaluated tip reuse performance under the bleach-based cleaning workflow used in high-throughput turbidostat operation. Expected versus observed volumes were measured using a brand-new box of Festo-compatible tips and compared to tips that had undergone repeated bleach sterilization and reuse cycles. No systematic drift in delivered volume was detected after reuse (mean absolute error 4.14 uL reused vs 4.05 uL new; t = 0.11, p > 0.9; **(Fig. 5D)**, and error distributions remained comparable to those obtained with new tips. Together, these results demonstrate that the OLH exceeds manual pipetting accuracy with both fresh and repeatedly sterilized tips (RMSE 7.00 uL and 5.42 uL, respectively, vs 10.45 uL manual), meeting the dosing requirements necessary for stable, long-duration closed-loop culture control.

**Figure 5).**
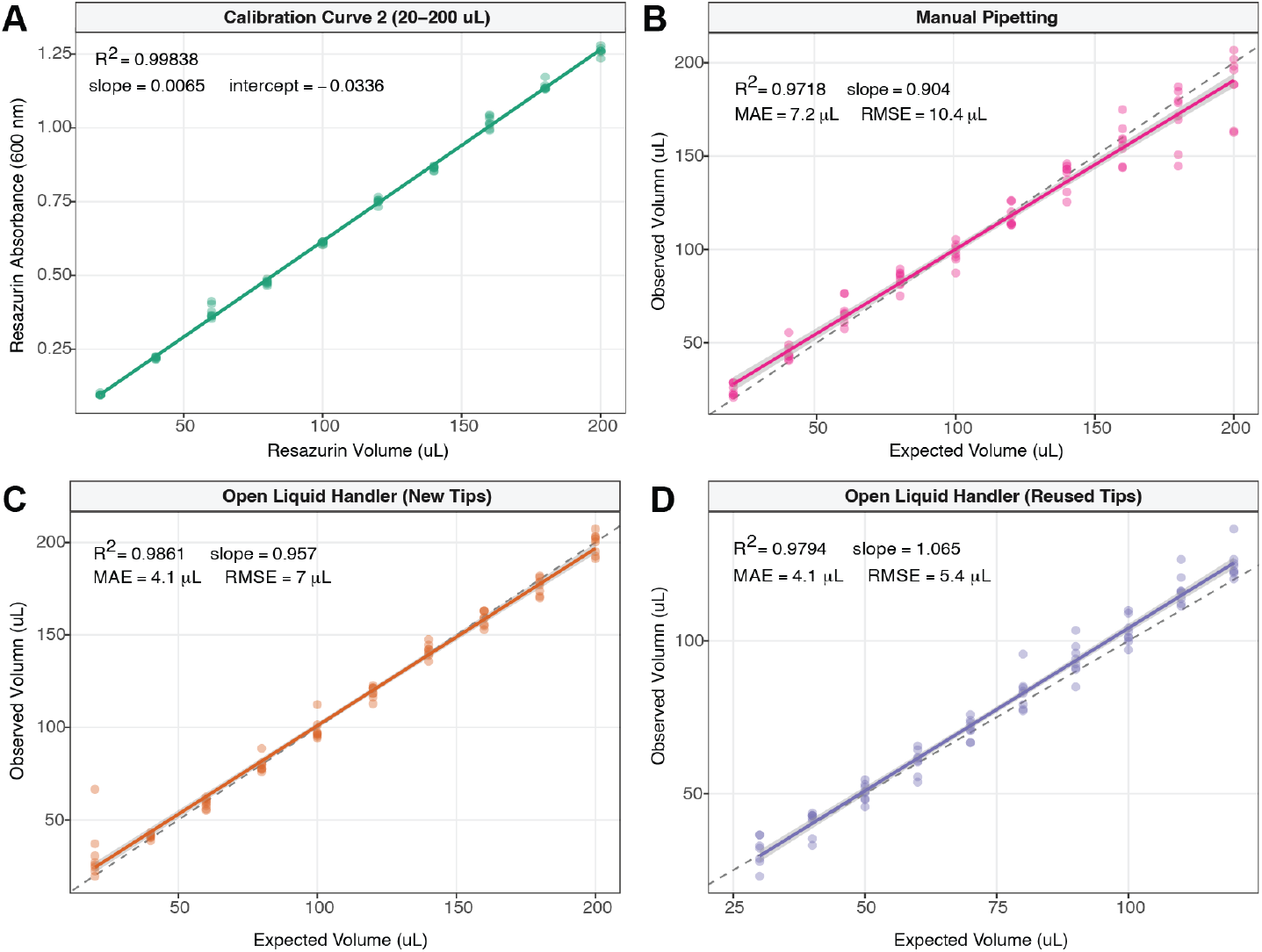
Pipetting accuracy validation for the Open Liquid Handler. **(A)** Absorbance-based calibration curve generated by manually pipetting serial dilutions of resazurin (20-200 uL, eight replicate wells per target volume) and measuring blank-subtracted absorbance at 600 nm on a Byonoy Absorbance 96 plate reader (R^2^ = 0.9984, slope = 0.0065 AU/uL, n = 79). This curve served as the reference standard for converting absorbance to effective volume in all subsequent panels. **(B)** Manual multi-channel pipetting with a P1000 (n = 80). **(C)** Open Liquid Handler with new tips (n = 80). (D) Open Liquid Handler with reused, bleach-sterilized tips (n = 79; volume range 30-120 uL). Each point represents a single well, solid lines show ordinary least-squares linear fits with 95% confidence intervals, and the dashed line indicates perfect agreement (slope = 1, intercept = 0). The OLH with new tips showed significantly lower mean absolute error than manual pipetting (MAE = 4.1 vs 7.2 uL; Welch’s t-test, t = -2.91, p < 0.005) and approximately half the error variance. Reused tips performed comparably to new tips (MAE=4.1 uL; t = 0.11, p > 0.9), indicating no systematic degradation from sterilization.

### High Throughput Turbidostat Implementation

High-throughput turbidostat control is a stringent case study for time-sensitive, closed-loop biological automation because timing is governed by biological constraints. In a prior implementation on a standard liquid-handling platform^7^, tip sterilization time is the dominant bottleneck. An 8-channel head performs bleach handling, and additional bleach and water rinses are parallelized by switching to a 96-channel head. Without the 96-channel head, the wash and rinse sequence becomes rate-limiting and the system cannot maintain the control cadences required for standard laboratory bacteria strains. The OLH is designed so that rapid point-to-point motion, short travel paths, and an integrated wash workflow support these cadences without relying on a second high-parallelism pipetting head. A standard turbidostat maintains culture density by coupling media exchange to turbidity measurements **(Fig. 6A)**. In the plate-based implementation, the workflow maintains microbial cultures near a defined optical density setpoint by repeatedly measuring culture density and applying per-well dilution decisions on a cadence matched to growth dynamics **(Fig. 6B)**. Density excursions can create confounds by shifting growth phase, resource availability, and effective selection pressure, and latency between measurement and actuation directly affects stability in long-running experiments. The method therefore uses a strict repeated control cycle that couples OD measurement to intervention with a short, end-to-end loop and is designed to scale to dozens to hundreds of cultures while preserving per-culture independence.

**Figure 6).**
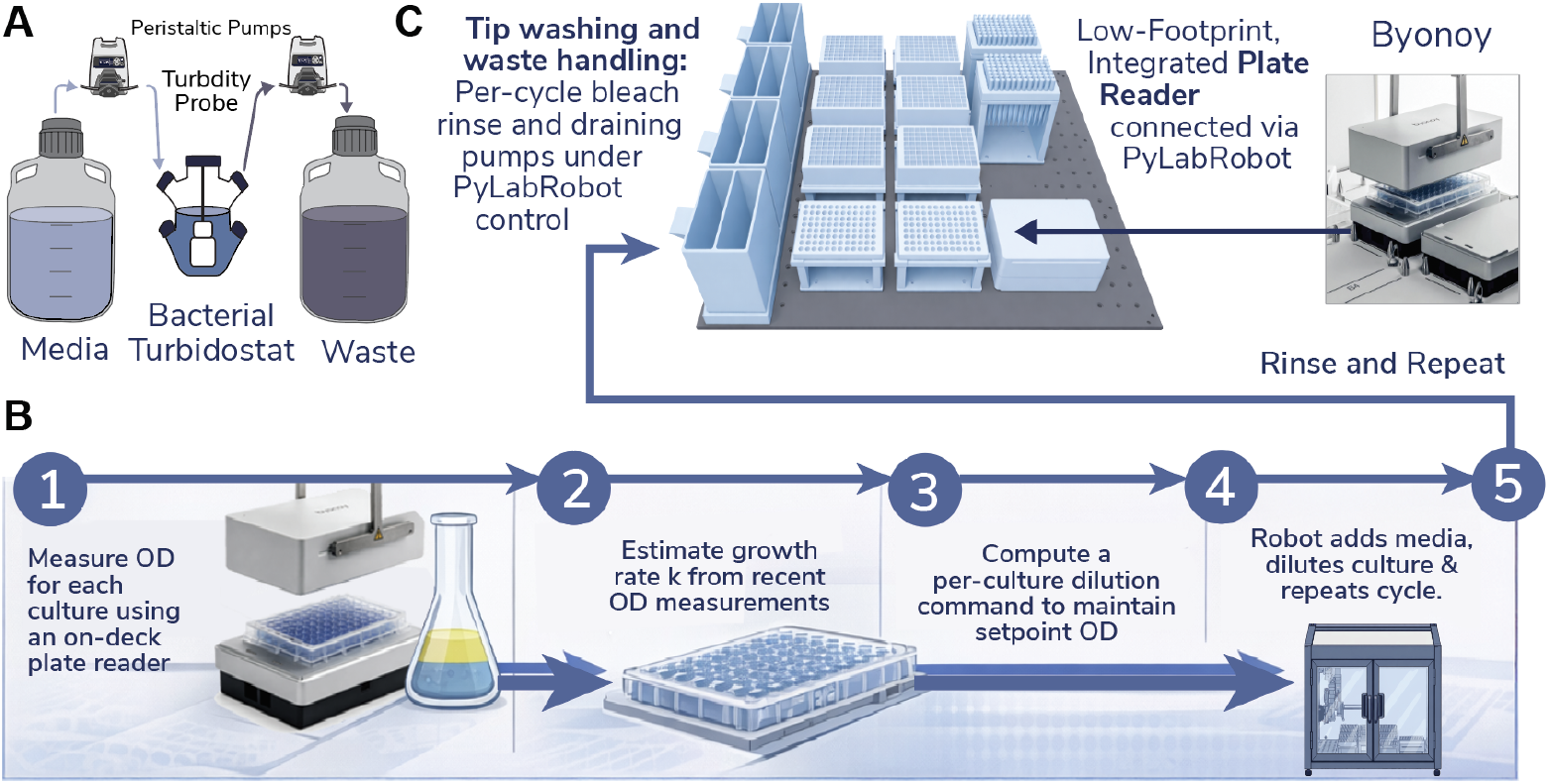
High-throughput turbidostat implementation on the OLH. **(A)** Standard turbidostat culture handling. Media and waste reservoirs are interfaced to a bacterial culture vessel, with fluid exchange driven by peristaltic pumping and turbidity or OD measurements used for feedback control. **(B)** Closed-loop control cycle executed by the OLH: (1) measure OD for each culture using an on-deck plate reader, (2) estimate per-culture growth rate from recent OD measurements, (3) compute a per-culture dilution command to maintain a target OD setpoint, and (4) execute media addition to dilute cultures, repeating the cycle at a fixed cadence. **(C)** Integrated tip washing and waste handling workflow under PyLabRobot control, including per-cycle bleach rinse and waste draining, and an enclosed, low-footprint plate reader (Byonoy Absorbance 96) that occupies a single deck position to minimize instrument footprint while maintaining rapid read-to-actuate timing.

The measurement step uses an on-deck Byonoy Absorbance 96 plate reader that occupies a single deck position and fits within the enclosed workspace **(Fig. 6C)**. This compact integration eliminates the need for a standalone, external plate reader or a dedicated plate-transport path, or transfer station into a separate instrument. Because the reader remains fixed on-deck, OD acquisition is scheduled as a short transfer and read operation within the same control loop, reducing latency between sensing and actuation while preserving deck capacity for tips, media, waste handling, and washing. Each cycle includes plate measurement, per-well decision-making, scheduled liquid handling execution, automated sanitation and waste handling, and logging and state update. OD is acquired as a full-plate matrix from the on-deck reader and is passed directly into the controller. Per-well dilution volumes are computed from the current OD value and recent OD history, using a growth estimate over the most recent interval when OD is in a reliable range. The controller outputs a media addition volume for each well along with safety and quality constraints, including skip logic for wells that do not require intervention and caps that limit maximum per-cycle dilution volume. The liquid handling step executes computed dilutions with an explicit scheduling strategy to preserve throughput, including batching to minimize travel overhead and skipping wells below intervention thresholds to reduce cycle time and consumable use. Automated washing and waste-path sterilization are integrated as first-class operations in the cycle, with bleach rinse and waste draining routines used to maintain long-run stability. Typical defaults used in the current implementation include an OD setpoint of 0.8, a working volume of 150 uL per well, a maximum dilution volume of 200 uL per cycle, and a nominal cycle interval of approximately 25 minutes. These values are each tunable per strain, reactor, and condition to balance growth dynamics with the time budget required for motion, reading, dispensing, and washing.

## Discussion

This work presents the first fully open, buildable liquid-handling platform assembled entirely from commercially available industrial motion, pneumatic, and control components, with a small number of machined or 3d printable adapter plates and deck fixtures. The instrument is operated through an open control stack integrated with PyLabRobot, so protocols and device integrations remain accessible in a single Python layer. A demanding closed-loop turbidostat workflow is used as a systems-level demonstration because it forces the core performance claims to be true: motion throughput, low-latency measurement, repeatable dosing, and reliable cleaning cannot be treated as secondary features when biology is time-sensitive.

This turbidostat case study directly reinforces why hardware customization is a needed feature in laboratory automation. In time-sensitive biology, control over kinematics and scheduling is not a luxury, it is the experiment. When service time per culture sets the control cadence, tunable motion profiles and short travel paths directly expand the range of growth rates that can be stabilized and the complexity of workflows that can be multiplexed. In practical terms, performance tuning converts “can the robot run this method” into “what region of biological dynamics becomes feasible under closed-loop control.”

These capabilities are particularly relevant for future applications including growth phenotyping of clinical isolates under antibiotic pressure, where phase-resolved, steady-state measurements are difficult to obtain with conventional batch handling. Density excursions, lag transitions, and measurement latency can blur phenotypes that only appear at specific growth states, and endpoint readouts such as Minimum Inhibitory Concentration (MIC) collapse rich dynamics into a single threshold. Automated closed-loop control supports longitudinal, quantitative phenotypes, including growth-rate shifts, adaptation dynamics, and recovery trajectories under time-varying perturbations. The same pattern generalizes to other domains where timing is the variable of interest, including directed evolution and adaptive laboratory evolution, combinatorial condition screening, and longitudinal perturbation experiments that require consistent, auditable intervention.

Several limitations remain and should be treated as guidance for users. Optical density measurements can be noisy or biased by bubbles, aggregation, meniscus effects, or path-length differences, which can propagate into control decisions if filtering and masking are not applied. Long runtimes and repeated liquid handling increase contamination risk, so sanitation routines, tip handling rules, consumable reuse limits, and waste sterilization are critical factors for success. Build reproducibility depends on assembly skill and rigorous calibration, and throughput can trade off against accuracy when aggressive motion profiles excite vibration or when pneumatic timing is pushed beyond validated regimes. Finally, some workflows will require different end effectors, sensors, or enclosure configurations, and the platform shown here should be viewed as a starting point for purpose-built variants rather than a universal substitute for every liquid-handling task.

Not surprisingly, development exposed predictable failure modes at the mechanical, software, and biological levels, and each category shaped the implementation. Mechanically, alignment, backlash, vibration, cable drag, and tip pickup reliability constrain both speed and repeatability, motivating explicit alignment checks, conservative cable routing, and calibration routines that can be repeated after service. In software, scheduling edge cases, error recovery, long-run logging, and state management become first-order requirements when experiments span days, which motivated resumable execution and an audit trail that ties measurements to actions. Biologically, evaporation, foaming, biofilm formation, clumping, and carryover can destabilize control; mitigation relies on containment, consistent washing and waste sterilization, and control logic that tolerates outliers rather than amplifying them.Near-term improvements fall into three categories. On the hardware side, tighter enclosure sealing, optional airflow management (for example HEPA-filtered inflow), more integrated waste handling, and improved end-effector modularity would reduce operational friction and broaden compatible assays; tool-changing and additional multi-channel options would expand throughput regimes without sacrificing task flexibility. On the control side, more advanced policies (for example adaptive filtering, model-based estimation, or model predictive control) and automatic anomaly detection could improve stability under measurement artifacts and heterogeneous growth. On the software and developer side, standardized configuration files, stronger simulation support, protocol validation tools, and expanded peripheral drivers would lower the barrier for new users while improving reproducibility across sites.

Overall, our long-term goal is to lay the groundwork for a modular, user and community-driven automation platform, where the build, control software, and documentation are designed to be extended like Lego blocks as new features and workflows are added. By completing a full replica build using only the bill of materials and compiled assembly guide (with two lab members spanning ∼1.5 weeks), we demonstrate that the system is fully reproducible and readily deployable. That said, the replica process also surfaced practical issues that were not apparent during the initial build (for example, motor cable routing requires manual finesse and planning to ensure sufficient slack is distributed in the cables), and resolving these issues continue to inform the assembly guide, troubleshooting documentation, and recommended commissioning tests. In practice, successful assembly from the instructions provided can be accomplished by someone skilled in mechanical and electrical assembly, but does not require formal mechanical or electrical engineering education as a prerequisite.

Near-term, we plan to expand demonstrations to additional closed-loop culture and screening workflows, but the core idea reaches far beyond bacterial growth control. An open-source control stack that bridges the gap from data analysis to low-level hardware control supports many workflows where measurements drive real-time experimental planning and decision-making, including adaptive cell culture and organoid maintenance^17^, differentiation and stimulation protocols in mammalian systems, dynamic formulation and reaction optimization in chemistry and materials synthesis^17,18^, iterative gene library construction and screening^19^, sample processing pipelines that branch based on QC metrics, and instrument orchestration for imaging, sequencing preparation, or analytical assays^20^. The capability of an open liquid-handling platform to seamlessly integrate with sensor and measurement data is uniquely advantageous for integration with self-driving labs and AI-guided decision making, which have been receiving increasing interest from both industry and academia across many fields of discovery^21–24^. More broadly, self-built, feedback-responsive robots make it possible to move from static protocols to adaptive experimentation, enabling compact, contained systems that are open to community extension, and can be specialized for diverse environments, constraints, and applications.

Because all components are commercially available and the build does not depend on specialized facilities, the platform can, in principle, be deployed rapidly and in parallel across multiple sites, a property that is especially relevant for surge capacity during public health emergencies or for distributed campaigns where purpose-built instruments are needed faster than commercial lead times allow. As a result, labs can compare methods directly and adapt the platform to their constraints rather than treating automation as a static appliance. In that context, the methods and systems presented here are a starting point: a complete reference implementation that others can reproduce, modify, and build on to engineer a broader suite of purpose-built, data-responsive robots for emerging scientific needs.

## Acknowledgements

This project was developed in collaboration with members of the Festo Corporation, who generously provided materials and technical components to enable the build and testing described here. We thank the PyLabRobot community for continued support and development, and Byonoy for the donation of the plate reader for development purposes. This work was additionally supported through generous funding to the Chory Lab by Duke University as well as an award from the Hypothesis Fund to E.J.C. Additionally, work was made possible with support through the National Institute of General Medical Sciences funded, Biomolecular and Tissue Engineering Training grant awarded to S.M.G. (T32-GM144291).

## Code and build package availability

The PyLabRobot turbidostat control code is currently available on GitHub^25^. Upon final publication, we will release a complete open build package designed to enable full replication and modification of the OLH platform. This package will include mechanical CAD files and 3D-printable part models, a detailed bill of materials with catalog numbers and supplier information, wiring diagrams and I/O mapping tables for all electrical and control components, step-by-step assembly and calibration documentation, and the full source code for hardware control, experiment scheduling, and data analysis. We are finalizing this documentation for public release and it is not yet posted in full due to the scope of the materials involved. For status updates or early access to in-progress documentation, contact.

## Methods

### Calibration Curve Generation

An absorbance-based standard curve was generated using serial dilutions of resazurin dye to establish a quantitative relationship between absorbance and dispensed volume. Resazurin is commonly quantified by fluorescence (excitation 530-570 nm, emission 580-620 nm), but can also be measured colorimetrically at 600 nm. Here, we used absorbance detection at 600 nm on a BMG Vantastar plate reader. Although absorbance-based detection of resazurin is less sensitive than fluorescence, it was sufficient for the volume range tested and allowed all calibration and validation measurements to be performed on the instrument integrated into the OLH deck. Resazurin stock solution was prepared at a known concentration and diluted in a 96-well microplate across a defined volume range (20-200 uL in 20 uL increments) using a calibrated manual P200 pipette. Eight replicate wells (rows A-H) were prepared at each target volume (n= 80 total observations). A linear regression of absorbance versus expected volume was fit to derive a transfer function for converting absorbance to effective volume. This single calibration curve was used as the reference standard for all subsequent pipetting validation experiments.

### Pipetting Accuracy Validation

To benchmark volumetric accuracy, three pipetting conditions were evaluated against the calibration curve: (1) the Open Liquid Handler with new tips (“Liquid Handler (New Tips)”), (2) manual multi-channel pipetting with a P1000 (“Manual Pipetting”), and (3) the Open Liquid Handler with bleach-sterilized reused tips (“Liquid Handler (Reused Tips)”). For conditions 1 and 2, serial dilutions spanning 20-200 uL in 20 uL increments were dispensed into 96-well plates with eight replicates per target volume (n = 80 observations). All eight channels of the OLH pipetting head (channels 0-7, corresponding to rows H-A) were used. For condition 3, dilutions spanning 30-120 uL in 10 uL increments were dispensed across all eight channels (n = 80 observations). OD600 was measured on the same plate reader used for calibration, and absorbance values were converted to effective volumes using the calibration transfer function. Volumetric error was computed as the difference between measured and expected volume for each well.

### Statistical Analysis

All statistical analyses were performed in R using the ggplot2, dplyr, tidyr, and readr packages. For each condition, ordinary least-squares linear regression was fit to measured versus expected volume to obtain the coefficient of determination (R^2^), slope, and intercept. Accuracy metrics included mean absolute error (MAE) and root mean squared error (RMSE) across all volumes tested. To compare OLH and manual pipetting accuracy, a one-sided Welch’s t-test was applied to the absolute errors (H_0_: OLH absolute error ≥ manual; H_1_: OLH absolute error < manual). A Wilcoxon rank-sum test was also performed as a nonparametric robustness check. Precision was assessed by comparing error variances between conditions using an F-test. To evaluate whether tip reuse affected pipetting performance, a two-sided Welch’s t-test was applied to absolute errors between the new-tip and reused-tip conditions. For visualization, expected versus measured volume was plotted for all four conditions (calibration curve, manual pipetting, liquid handler with new tips, liquid handler with reused tips) in a 2 × 2 faceted panel using ggplot2 with geom_smooth (method = “lm”) and 95% confidence intervals. Per-panel annotations include R^2^, slope, MAE, RMSE, and sample size. A dashed identity line (slope = 1, intercept = 0) is shown for reference. All raw data from both experiments were combined into a single annotated CSV file with standardized condition labels, computed percent error, and absolute error for reproducibility

**Supplemental Figure 1).**
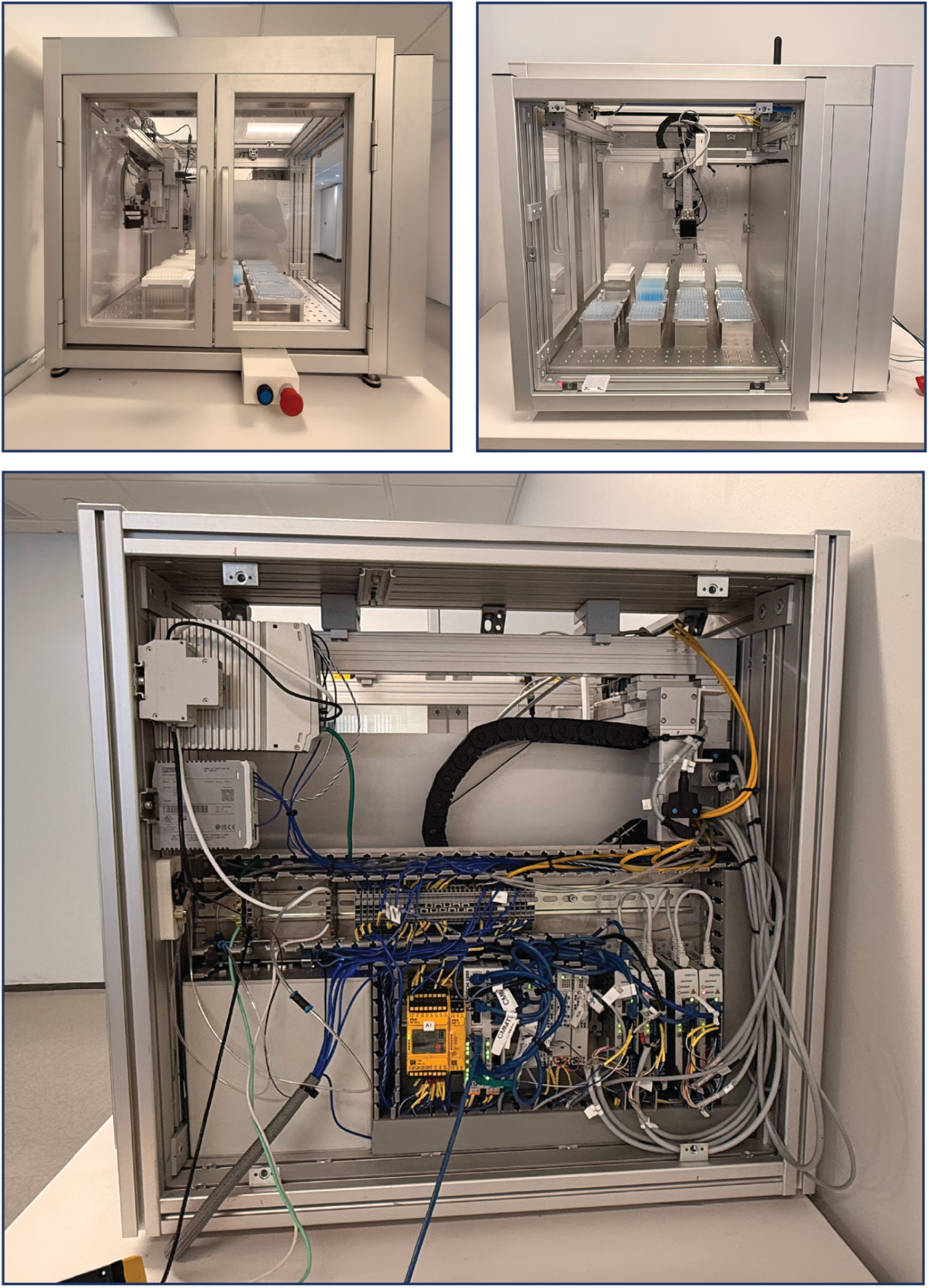
A Replica Build of the Open Liquid Hander. (top left) front view. (top right) side view. (bottom) back view of control cabinet. *Image backgrounds were digitally removed using Adobe Photoshop Generative Fill (AI-assisted editing) to reduce visual clutter; no instrument features were added, removed, or altered*.

**Supplemental Figure 2).**
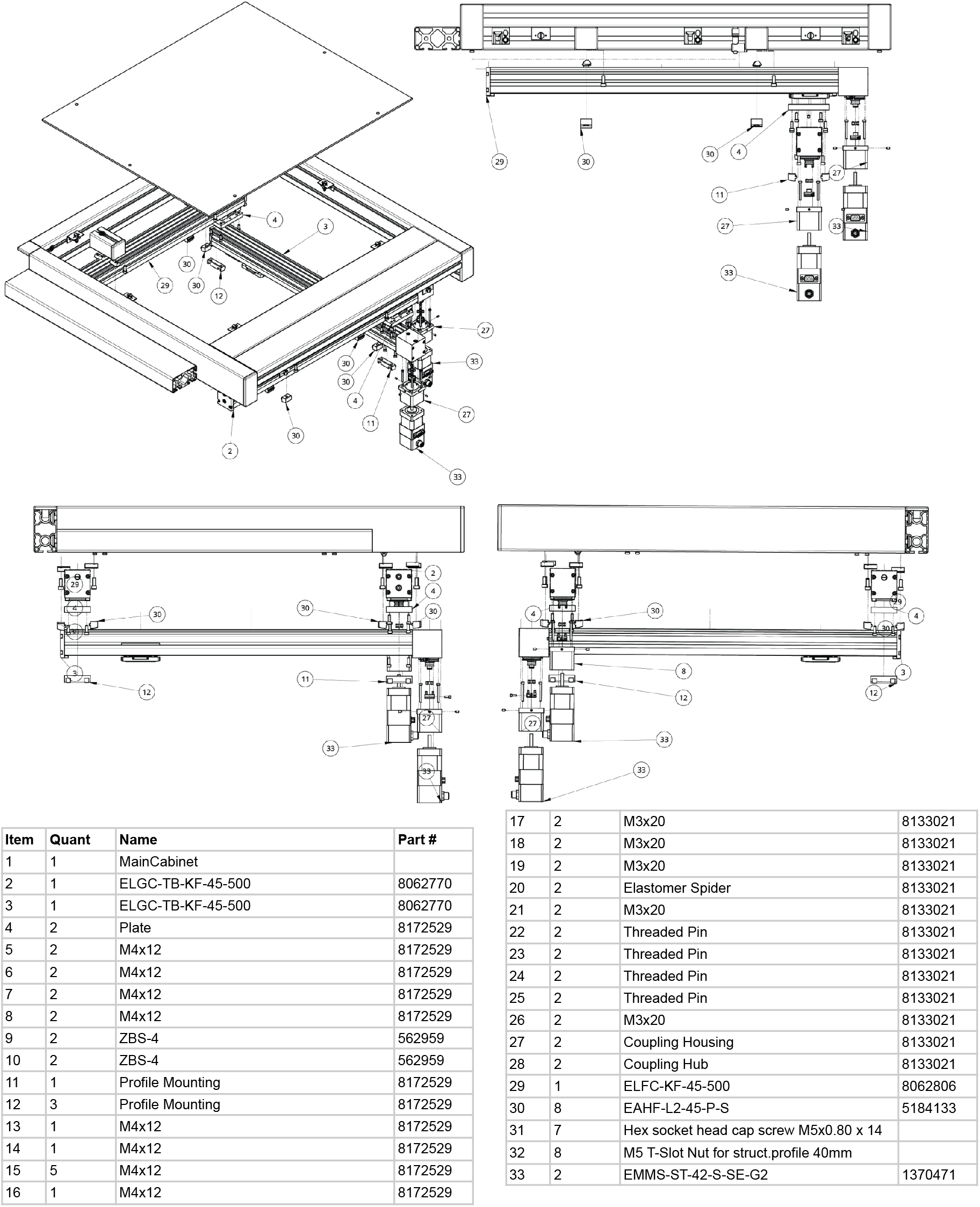
Gantry Assembly and views of the OLH motion stack. Isometric and orthogonal views show the X and Y linear axes, dual Z stages for pipetting and gripper modules, and mounting interfaces within the enclosure frame. Callouts correspond to bill of materials items listed below.

**Supplemental Figure 3).**
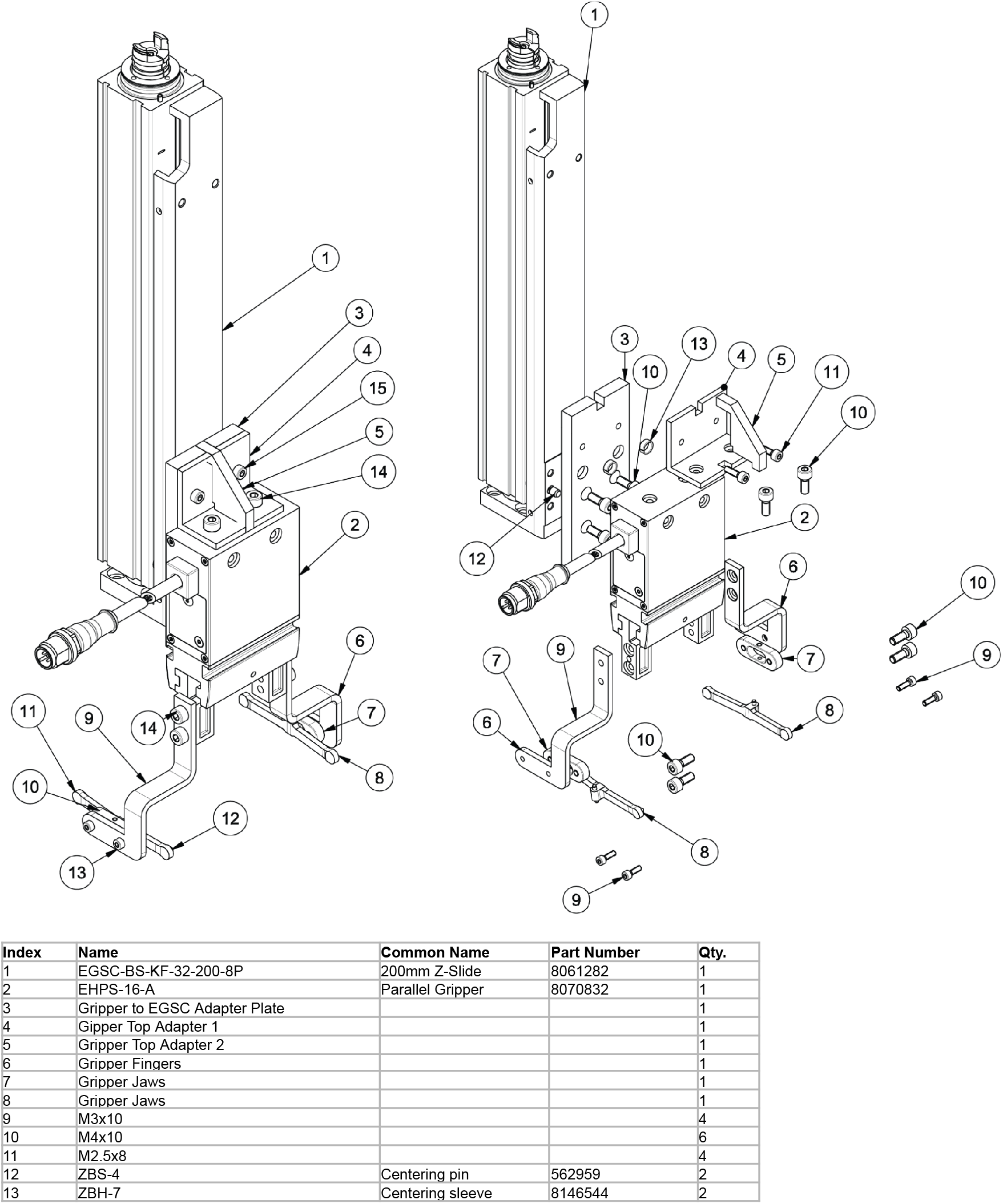
Gripper assembly. The parallel gripper is mounted to a 200 mm Z-slide and coupled through a linkage and bracket assembly for plate handling. Exploded components and callouts correspond to bill of materials items listed below.

**Supplemental Figure 4).**
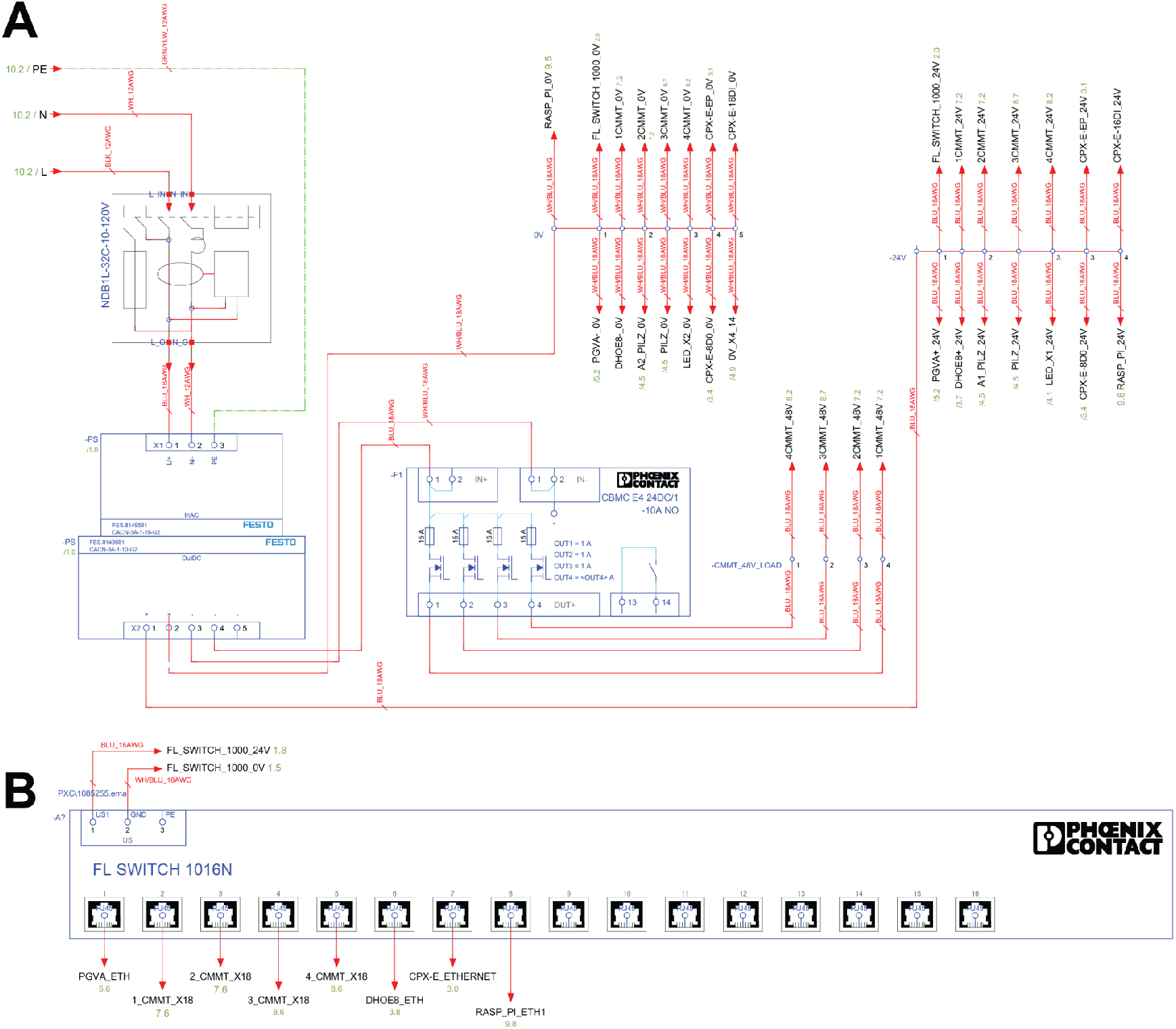
Power and networking architecture. **(A)** AC mains entry and 24 V DC power supply with protected distribution of 24V and 0V rails to cabinet subsystems. **(B)** Internal Ethernet switch wiring and network topology interconnecting the controller, motion drives, I/O, and peripherals.

**Supplemental Figure 5).**
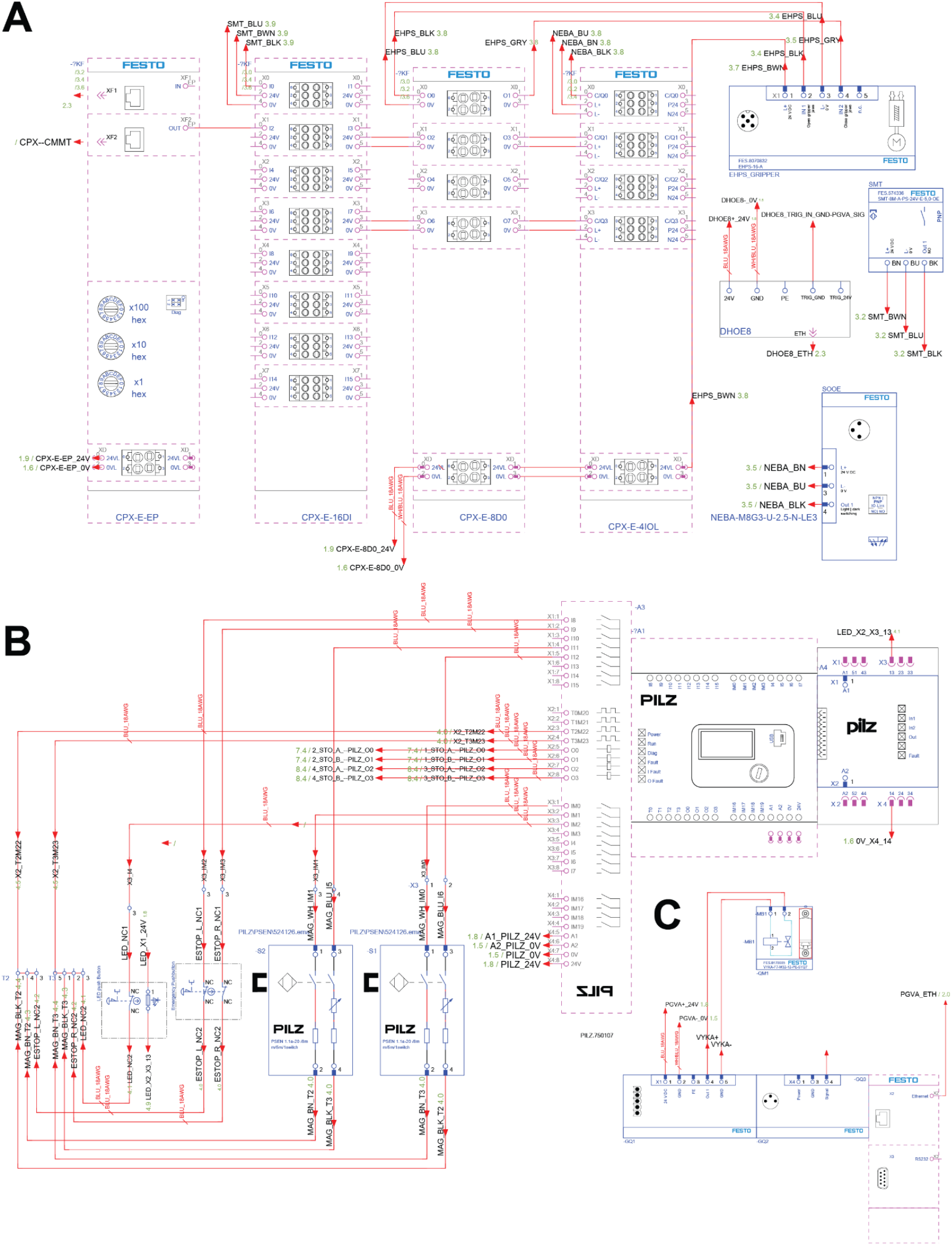
Control and safety architecture. **(A)** Distributed I/O system showing sensor and actuator wiring through the cabinet I/O modules. **(B)** Safety controller wiring with dual channel emergency stop inputs and motion disable outputs. **(C)** Peripheral control module wiring with cabinet power and communication interfaces.

**Supplemental Figure 6).**
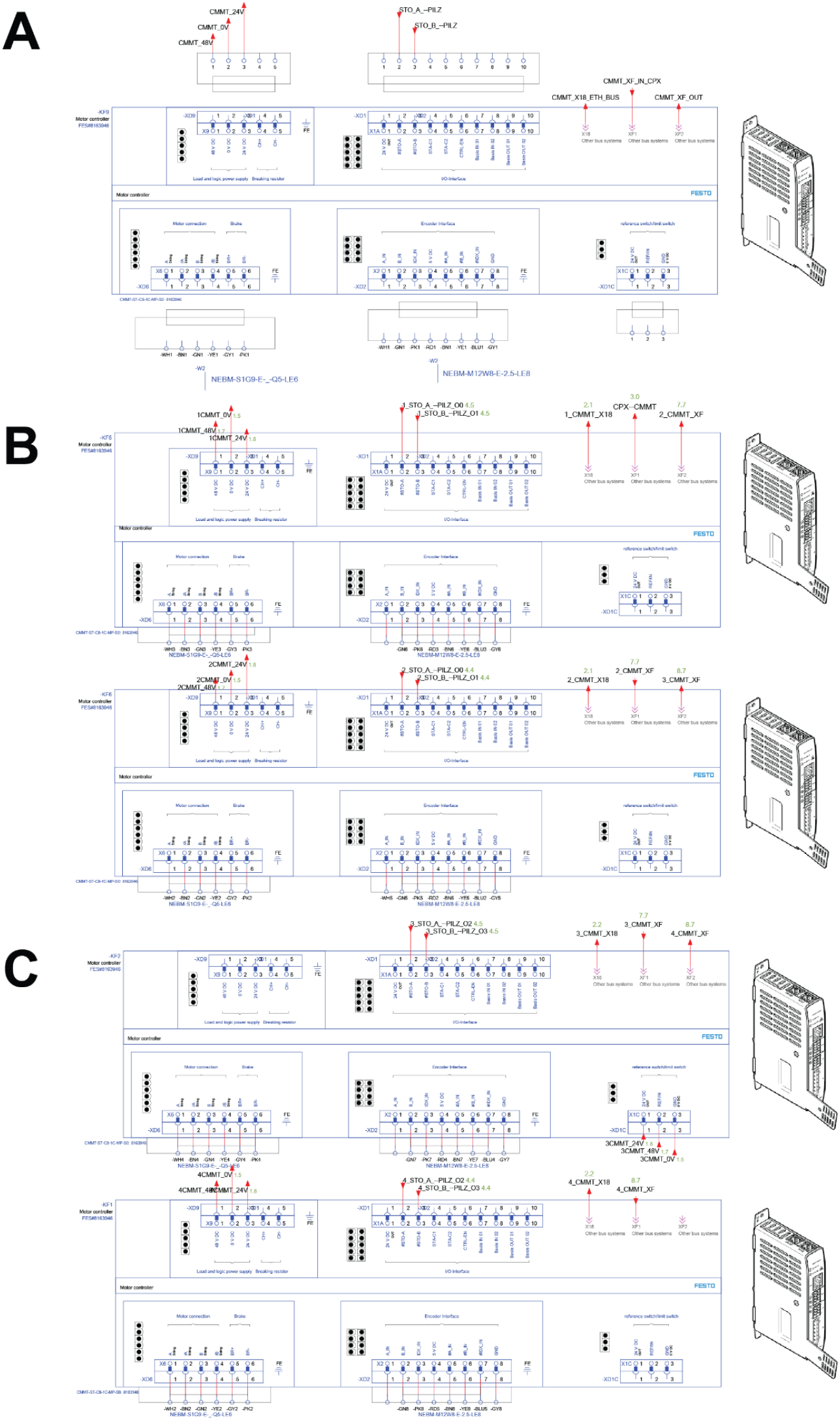
Motor drive architecture. **(A)** Motor drive wiring for the first axis, including power inputs, safety disable inputs, and motor and encoder connections. **(B)** Motor drive wiring for additional axes, showing replicated power, safety, and feedback interfaces. **(C)** Motor drive wiring completing the remaining axes with identical integration of power, safety, and feedback connections.

**Supplemental Figure 7).**
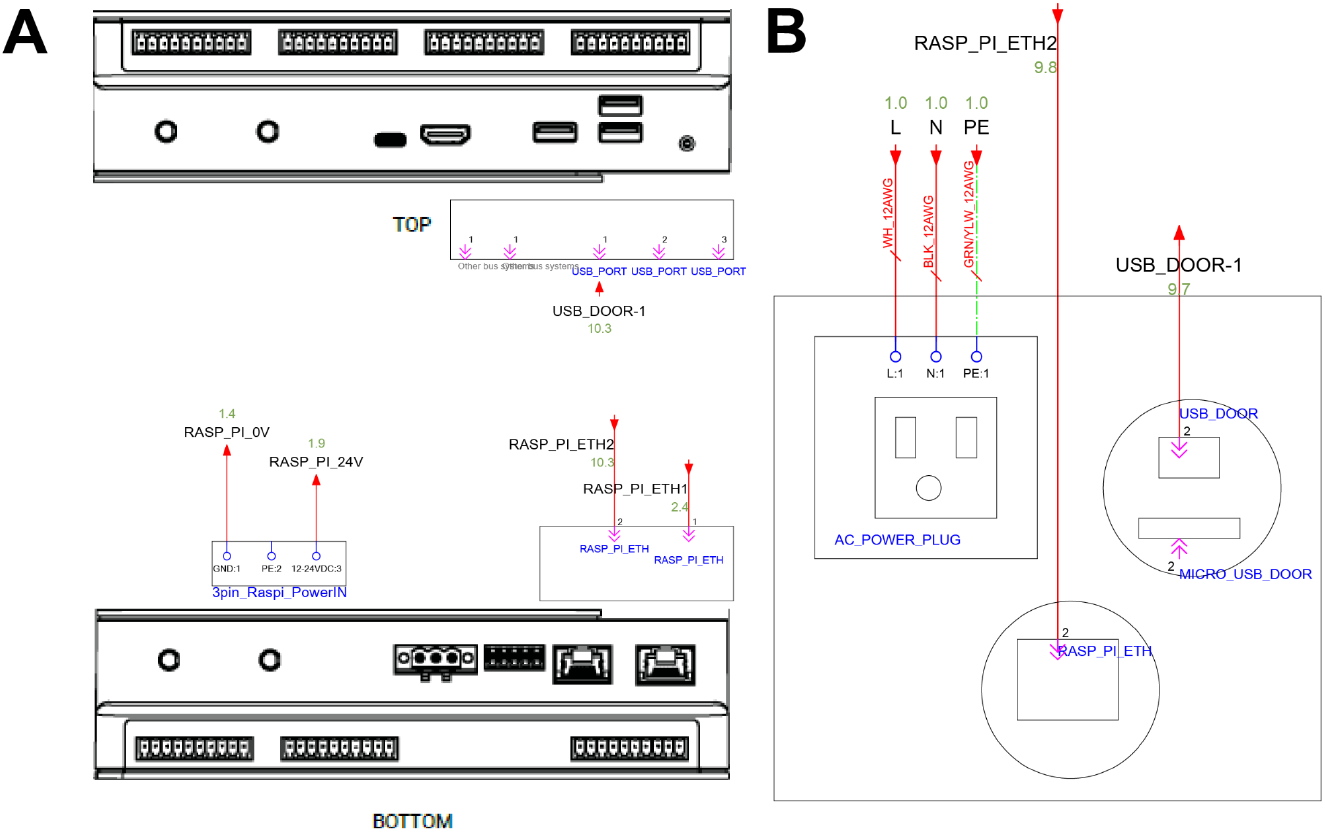
Supervisory compute and external interfaces. **(A)** Embedded computer wiring, including DC power input, Ethernet connectivity, and USB interfaces. **(B)** Door mounted service ports, including AC inlet and external data pass through connections.

**Supplemental Figure 8).**
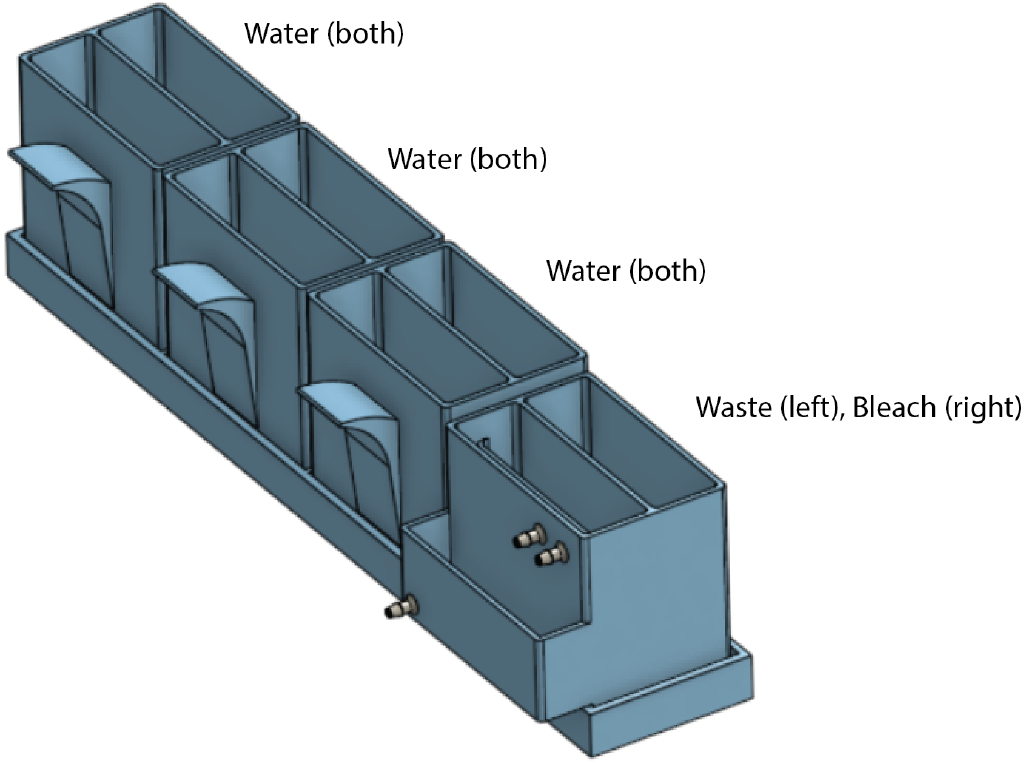
3D Printed Washer. The 3D printed washer has 3 2-compartment static reservoirs and a reservoir with 2 refillable compartments. One compartment on the refillable reservoir is used to collect and drain waste, and the other is to act as a bleach reservoir for sterilizing tips. Refillable reservoirs are connected to Agrowtek pumps through 3/16” tubing attached to barbed channel inlets on the washers. Static reservoirs are used for sequential water rinses following tip sterilization.

## Notes

### Competing Interest Statement

The authors have declared no competing interest.

